# Multi-modal characterization of rodent tooth development

**DOI:** 10.1101/2024.11.01.621612

**Authors:** Yuchen Jiang, Kaitlin A. Katsura, Nir Z. Badt, Marius Didziokas, Sonia Dougherty, David L. Goldsby, Elizabeth J. Bhoj, Kyle Vining

## Abstract

Craniofacial tissues undergo hard tissue development through mineralization and changes in physicochemical properties. This study investigates the mechanical and chemical properties of developing enamel, dentin, and bone in the mouse mandible. We employ a multi-modal, multi-scale analysis of the developing incisor and first molar at postnatal day 12 by integrating micro-computed tomography (microCT), nanoindentation (NI), energy dispersive spectroscopy (EDS), and Raman spectroscopy. Our findings demonstrate distinct patterns of mechanical, elemental, and chemical changes across mineralized tissues. These results suggest that mineral composition drives mechanical properties across different craniofacial hard tissues. Integrating multi-modal characterization of mineralized tissues opens new opportunities for investigating structure-function relationships in craniofacial biology and genetics.

## Introduction

The development of mineralizing tissues is regulated by specialized groups of cells that undergo complex processes of formation and differentiation (1-3). While significant strides have been made in understanding the genetic (4, 5) and molecular mechanisms (6) of mineralization, a critical gap remains in understanding how these processes drive the physical properties of the surrounding tissues or matrices. One promising tool for studying the development of mineralized tissues is the rodent incisor, as it is continuously growing with a stem cell niche proliferating to form an ever-growing mineralized tip that gets abraded when eating (7, 8). Targeted genetic manipulations in mice have also demonstrated the mouse to be useful at dissecting the mechanisms for tooth-related diseases that affect mineral quality (Amelogenesis imperfecta) (9, 10) and tooth morphology (11), using a variety of immunohistochemical techniques. However, these methods primarily assess the cellular drivers of the matrix, whereas the extracellular matrix and minerals have been found to be critical drivers of cell expression. By integrating mechanical and elemental analysis of dental tissues, we can better understand the interplay between function and composition of tissues.

Teeth are composed of two major hard tissues, (1) dentin that surrounds the nerve and blood vessels, and (2) enamel that forms the outer tooth crown and is the exposed mineral visible in the mouth (12). In the developing tooth, a bony ridge called the alveolus covers the tooth prior to the tooth erupting in the mouth. In all teeth, the enamel and dentin both form a protein-rich matrix of mostly acerogenins and collagens, respectively, to lay a framework or scaffold for the future mineral to harden and replace (13). This occurs in all human teeth and mouse molars. In mice, however, the incisor features all stages of tooth development because mice incisors continuously develop at all stages, making them an excellent model system (14).

Previous studies on the hard tissue matrices have performed nanoindentation (NI), scanning electron microscopy (SEM), energy dispersive spectroscopy (EDS), and Raman spectroscopy on extracted permanent teeth from humans and genetic mouse models. However, there remains a lack of studies on the developing tooth. SEM has been extensively used to study the various natural forms of hard mineralized structures, including crustaceans (15, 16), mollusks (17, 18), rocks (19), teeth from fish (20, 21) and other mammals (22, 23). It has also been applied to human and rodent hard tissues (bone, enamel, dentin) (24-30) to analyze the ultrastructures that define mineral quality and crystal architectures in the context of human disease; however, limitations exist due to the extensive preparation process. EDS and Raman (31, 32) have been performed to a lesser extent. While NI is an effective tool for studying rocks in geological settings, very limited work has been done on this precision mechanical testing method for developing hard tissues (33, 34). All these tools are powerful for studying mineralized substances, but they have yet to be integrated together to study the developing mouse incisor.

The developing tooth is a mineralizing composite that offers insights into the structure-function relationships. Here, we perform multi-modal materials characterization to mouse incisors to investigate the properties of the developing enamel and dentin. Our study builds on recent work showing that the molars of 12-week-old male mice can landmark the developmental stages of the continuously growing incisor (35) and an integrated method for tendon (36) and bone (37). We apply this research to earlier stages of post-natal development at day 12 (P12) in both female and male mice as a model. At P12, molar and incisor development are still underway, providing a critical window into early mineralization processes. By combining NI, EDS, and Raman spectroscopy from the same individual sample, we employ a multi-modal approach to study the mechanical and elemental properties of the craniofacial hard tissues. We hypothesized that mineral composition requires different elements at specific stages to contribute to hard tissue hardness rather than developing in a linear fashion.

In this paper, we developed an integrated framework for assessing how chemical composition relates to the hardness and modulus of dental tissues. By doing so, we identified the relation of enamel and dentin composition to their mechanical properties, as well as to craniofacial bone. By employing these engineering approaches in a developmental biology context, we can further expand our knowledge of the nuanced development in dental and craniofacial hard tissues.

## Results

### Mineralized density of developing mouse dentition peaks preceding eruption

Micro-computed tomography (microCT) imaging of whole mouse skulls at post-natal day 12 (P12) provided a comprehensive 3-dimensional view of the mineralization status in the developing incisor and molar. At this developmental stage, both the first molars and the continuously growing incisors exhibited comparable mineralized states, suggesting coordinated timing in their mineralization processes (Fig. 2). The ever-growing incisors showed that mineralization is initiated early and accelerates prior to eruption, in line with previous reports. Dentin mineralization commenced almost immediately, with continuous mineral deposition throughout development. Enamel mineralization, in contrast, follows a distinct spatial and temporal pattern. It initiated at the mesial region of the first molar (M1) and rapidly progressed distally along the incisor, reaching peak density just before eruption (Fig. 2). This peak mineralization coincided with the moment the incisor begins to erupt through the alveolar bone. Interestingly, the first molar followed a similar trajectory. However, it represented more of a snapshot of dentin and enamel development. The mineralization levels in the molar corresponded most similarly to that of the panel in Figure 2D, representing the developmental phase just before the incisor erupts. This alignment suggests that molar mineralization, while not continuous post-eruption like in incisors, achieved its peak density just prior to eruption. These findings highlight the dynamic and tissue-specific nature of mineralization across different tooth types, with incisors and molars following distinct yet synchronized mineralization timelines leading up to eruption.

**Figure 1.**
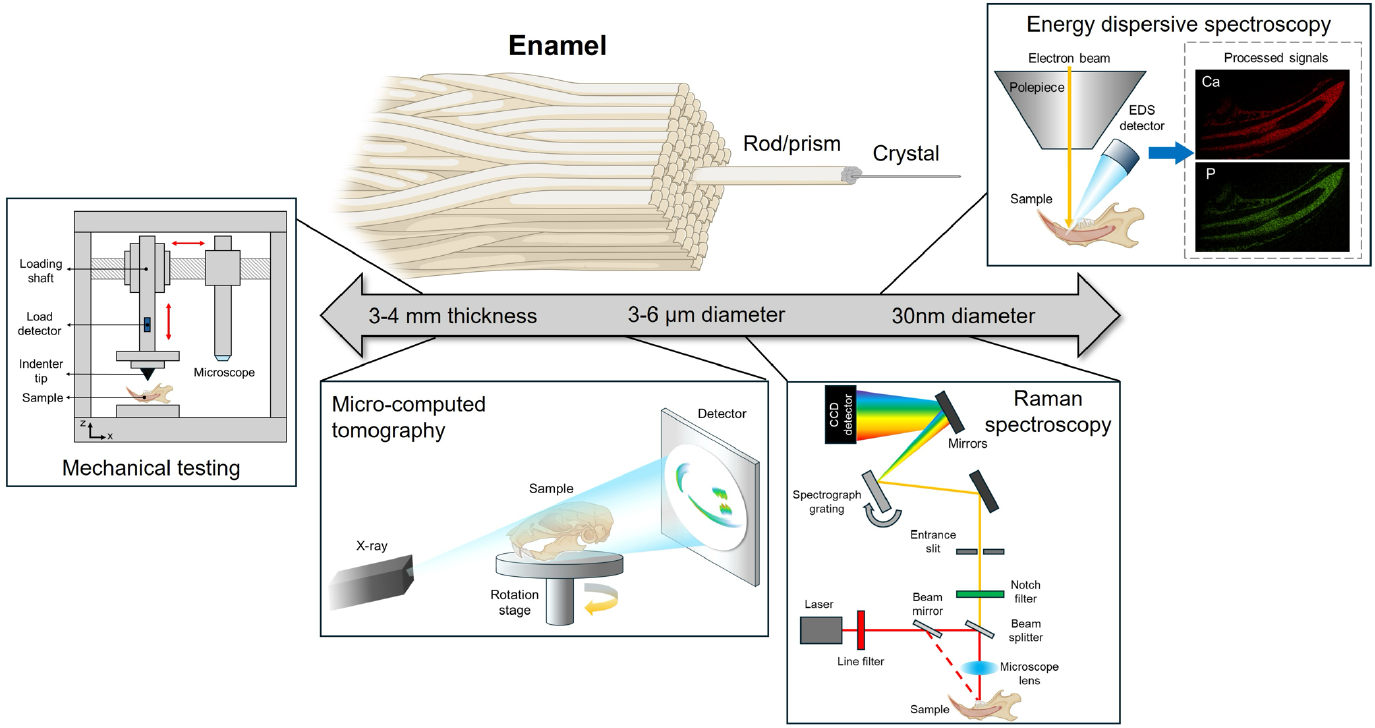
Outline of the multimodal methodologies used in this study across varying length scales. Here, enamel is used as the primary example due to its layered crystalline organization.

**Figure 2.**
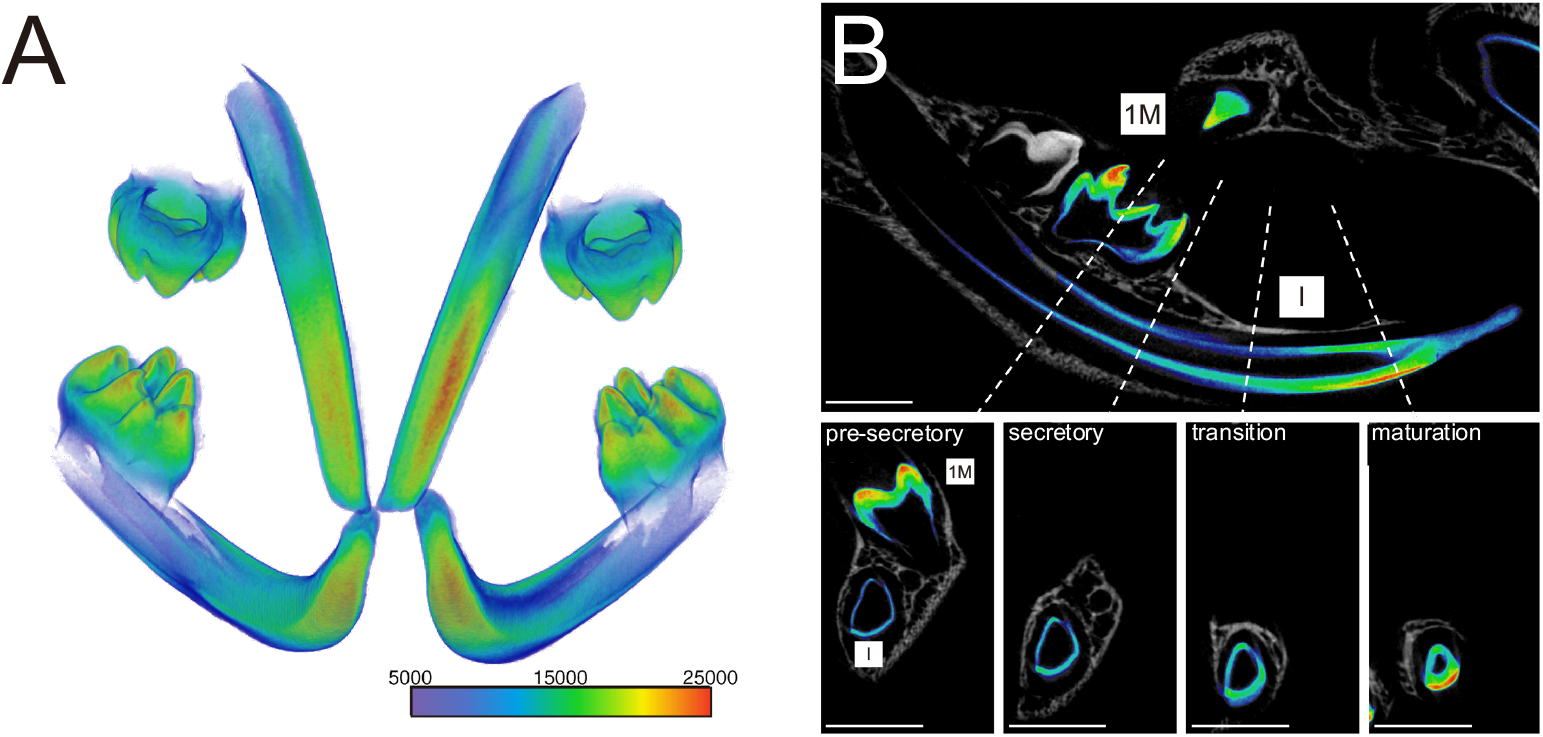
Micro-computed tomography (microCT) shows a 3-dimensional reconstruction of mineralizing dentition. (A) Mineralization heat map of the maxillary and mandibular incisors and molars showing the highest levels of mineralization as measured by greyscale concentrated at the molar cusp tips and incisor tips. (B) A 2-dimensional section through the sagittal plane of the mandible reveals a staged development of the ever-growing incisor and a first molar just prior to eruption from the alveolar bone. Labeled hashed lines (a-d) represent the cross-sectional views (below) at various stages of growth along the developing incisor, landmarked for pre-secretory stage (a), secretory (b), transition (c), and maturation (d) stages of enamel formation. Scale bars, 1mm.

### Mechanical property and elemental compositions of developing enamel diverge during mineralization

Paired nanoindentation and elemental analyses were performed on the incisor, molar, and bone of the same sample to explore the relationship between tooth hardness and elemental composition. Six indentations (marked by dashed triangles in Fig. 3A) were conducted to measure hardness on enamel (EM) and dentin (DT) across different developmental stages in both incisors and molars. Additionally, energy dispersive spectroscopy (EDS) line scans (shown as a yellow line in Fig. 3A) were conducted along the dentin-enamel junction at the same locations as the nanoindentation.

**Figure 3.**
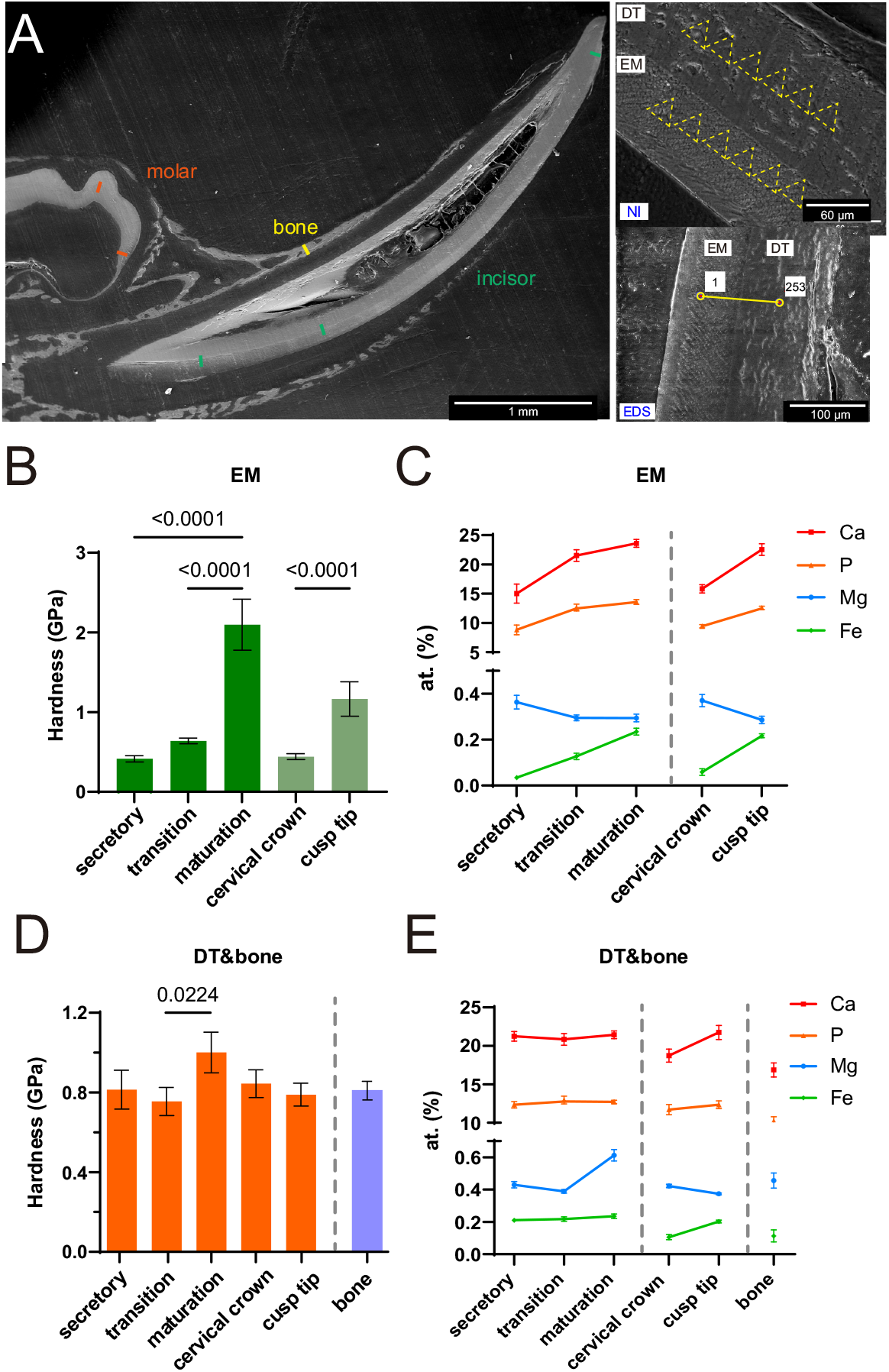
Nanoindentation and elemental analysis of rodent tooth development. (A) SEM image of a rodent incisor cross-section. Red marks indicate measurement regions, and the green marker highlights the alveolar bone near the first molar. Insets show six nanoindentation sites (dashed triangles) on enamel (EM) and dentin (DT) and an EDS line scan (yellow line) across the enamel-dentin junction. (B) Hardness of enamel in incisor and molar at different developmental stages, showing a notable increase during tooth development. (C) EDS results for elemental composition (Ca, P, Fe, Mg) in enamel during mineralization in both incisor and molar. (D) Hardness of dentin at different stages and of alveolar bone, showing similar values between the cervical crown and alveolar bone. (E) Elemental trends in dentin show stable Ca, P, and Fe, with Mg influencing hardness during both dentin and alveolar bone mineralization.

In the incisor, enamel hardness increases modestly from the secretory to the transition stage, followed by a significant increase during maturation (p < 0.0001) (Fig. 3B). A similar trend was observed in molar enamel, where hardness increased markedly from the cervical crown to the cusp tip (p < 0.0001) (Fig. 3B). EDS analysis revealed notable variations in the atomic percentages of iron (Fe), calcium (Ca), magnesium (Mg), and phosphorus (P) during mineralization in both incisors and molars. In enamel, Ca and P levels are substantially higher compared to Mg and Fe, consistent with previous studies (38-40). Across both incisor and molar samples, the atomic percentages of Ca, P, and Fe increase progressively through mineralization, while Mg shows an inverse trend—atomic percent is higher in the secretory stage and cervical crown but declines as mineralization proceeds (Fig. 3C).

In contrast, dentin hardness does not show significant differences between developmental stages, except for a relatively higher hardness during maturation (P=0.0224). The hardness of the alveolar bone, measured near the first molar (Fig. 3A), is comparable to that of the cusp tip (Fig. 3D). To correlate the mechanical data with elemental composition, EDS results indicate that the atomic percentages of Ca, P, and Fe remain relatively stable across the three developmental stages in the incisor, while Mg content decreases slightly from the secretory to transition stage and increases significantly during maturation (Fig. 3E). Interestingly, in dentin, Ca and Fe levels are lower in the cervical crown compared to the cusp tip, while Mg shows a slight decrease during molar development. Moreover, Ca, P, and Fe levels are lower in the alveolar bone compared to the cusp tip, with Mg content showing a similar increasing trend. These findings suggest that variations in elemental composition directly influence enamel and dentin hardness during tooth development. These findings suggest that Mg content in both dentin and alveolar bone plays a key role in influencing hardness during tooth and alveolar bone formation.

### Alterations of Chemical Composition During Tooth Mineralization

Raman spectroscopy was utilized to characterize the chemical compositions of developing incisors and molars at P12, building on previous applications of this technique on permanent teeth (41, 42). Measurements were taken at positions parallel to those analyzed by EDS and nanoindentation on the same sample, with three spectra collected per position to ensure reproducibility. Prominent Raman peaks were observed at 430 cm^−1^, 586 cm^−1^, 960 cm^−1^, and 1070 cm^−1^, corresponding to ν_2_(PO4^3−^), ν_4_(PO4^3−^), ν_1_(PO4^3−^), and ν_1_(CO3^2−^), respectively, which provide insights into the composition of mineralizing hydroxyapatite (Fig. 4A). Prominent Raman peaks were observed at 430 cm^−1^, 586 cm^−1^, 960 cm^−1^, and 1070 cm^−1^, corresponding to ν_2_(PO4^3−^), ν_4_(PO4^3−^), ν_1_(PO4^3−^), and ν_1_(CO3^2−^), respectively, which provide insights into the composition of mineralizing hydroxyapatite (Fig. 4A) (42-44).

**Figure 4.**
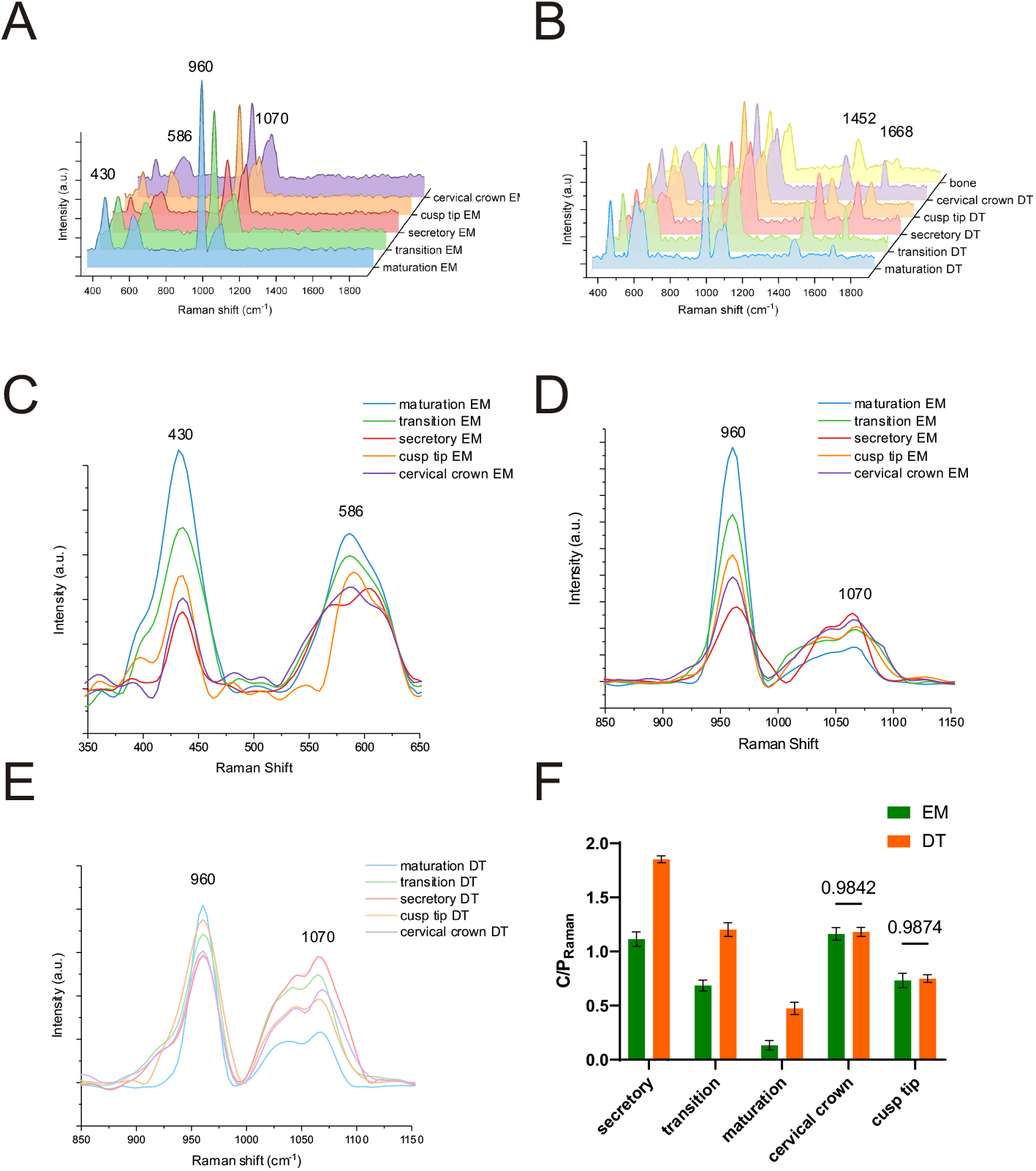
Raman spectra of (A) enamel and (B) dentin show varying levels of elemental compositions along incisor and molar mineralization. (C) The spectral region of the ν_2_, ν_4_ phosphate group, and (D) the spectral region of the ν1 phosphate and the ν_1_ b-type carbonate groups in enamel. (E) The carbonate-to-phosphate ratio (C/P) is represented as the ratio of the area under the 1070 cm^−1^ peak to the area under the 960 cm^−1^ peak.

Distinct variations in peak intensities and shifts were observed in both developing incisors and molars, reflecting different mineralization levels and organic content across regions. In enamel, phosphate bands at 430 cm^−1^ and 586 cm^−1^ displayed the highest intensities at the maturation stage, with a progressive decrease at the transition and secretory stages (Fig. 4C), and similar trend was also observed in dentin (SI Appendix, Fig. S1A). A similar trend was evident in molar enamel, where these bands exhibited higher intensities at the cusp tip compared to the cervical crown. Notably, the ν_4_ phosphate band showed an altered shape at the secretory stage, possibly due to variations in hydroxyapatite crystal orientation in this less mineralized region (45).

The ν_1_ carbonate and phosphate vibration modes exhibited prominent spectral changes in both enamel and dentin (Figs. 4A and 4B). In enamel, the intensity of the 960 cm^−1^ phosphate peak was markedly higher at the maturation stage and showed significant decreases from maturation to the secretory stage (Fig. 4D). By contrast, dentin displayed only minor intensity changes in both incisor and molar mineralization (Fig. 4E). Interestingly, the 1070 cm^−1^ carbonate band intensity decreased in both enamel and dentin as the mineralization of incisors and molars progressed.

To quantify these compositional changes, the carbonate-to-phosphate (C/P) ratio was calculated by comparing the area under the 1070 cm^−1^ carbonate peak to that of the 960 cm^−1^ phosphate peak (Fig. 4F). These band parameters were obtained by fitting Gaussian-Lorentzian curves in the 900–1130 cm^−1^ region (SI Appendix, Fig. S1B) (41, 43). Results indicated that the overall carbonate content in dentin was higher than in enamel along the incisor; however, no significant differences were observed between enamel and dentin in the molar (P = 0.9842 for cervical crown and P = 0.9874 for cusp tip) (Fig. 4F). Additionally, a clear decrease in the C/P ratio was observed in the incisor as mineralization advanced. Together, these findings show the distinct mineralization patterns of enamel and dentin and provide insights into the structural and compositional maturation processes in developing teeth.

### Integration of Mechanical and Compositional Properties using PCA and MLR

To integrate information from nanoindentation (NI), EDS, and Raman spectroscopy along the developing incisor and molar, we performed principal component analysis (PCA) with six continuous variables: NI-hardness, EDS/Mg (at%), EDS/Fe (at%), EDS/Ca (at%), EDS/P (at%), and the C/P ratio. PCA was used to reduce the dimensionality of the dataset while preserving variability by creating new, uncorrelated variables that maximize variance successively (46, 47) Principal Component 1 (PC1) accounted for 60.99% of the total variance (Eigenvalue 4.269), while Principal Component 2 (PC2) explained an additional 18.42% (Eigenvalue 1.290), with PC1 and PC2 collectively capturing 79.41% of the variance (SI Appendix, Table S1). PCA loadings indicated strong positive associations along PC1 for hardness and elemental concentrations of Ca, P, and Fe, whereas Mg and the C/P ratio were negatively associated with PC1. Notably, the C/P ratio diverged most strongly from hardness (Fig. 5A). Sample distribution along PC1 distinguished enamel developmental stages (secretory, transition, and maturation), with data points transitioning from positive to negative PC1 values. Conversely, tissue type was more influenced by PC2, with dentin clustering on the positive PC2 axis and enamel on the negative side. The loadings further suggest that EDS/Mg may play a key role in differentiating enamel from dentin, while other variables contribute more substantially to PC1 (38).

**Figure 5.**
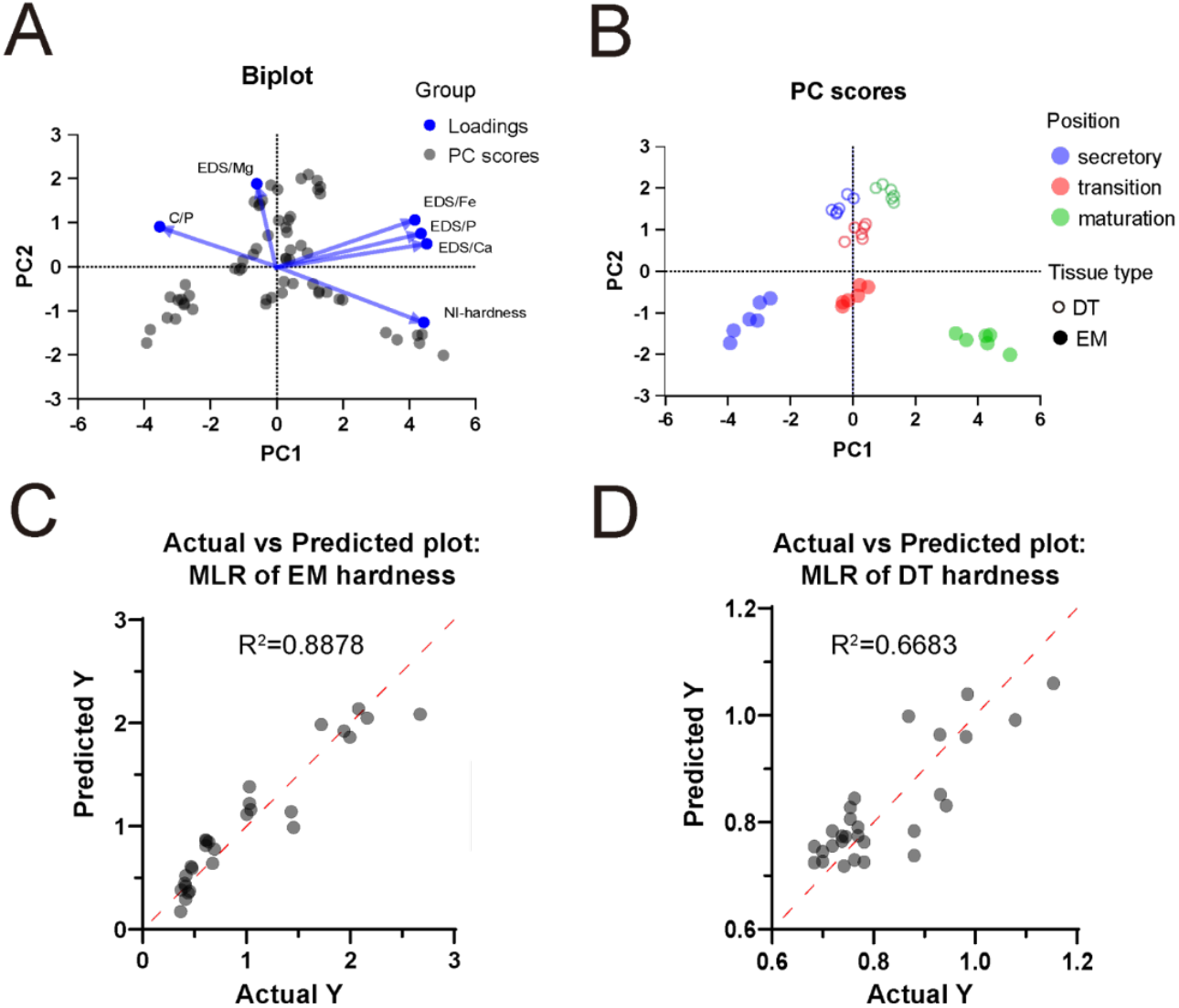
Principal component analysis (PCA) and multiple linear regression (MLR) integration models for enamel and dentin. (A) PCA biplot of enamel spectra illustrating sample clustering based on developmental stages, reflecting variations in elemental composition along the incisor and molar. (B) PCA scores show distinct clusters of enamel and dentin corresponding to different stages of mineralization along the incisor. (C) MLR model predicts hardness based on the input variables for (C) enamel and (D) dentin.

Examining enamel by tooth type (incisor vs. molar) revealed that the cervical portion of the molar clustered closely with the secretory stage of the incisor, while the cusp portion of the molar aligned with the incisor’s transition stage. The incisor’s maturation stage, however, was distinct. In contrast, dentin from both incisor and molar clustered together (SI Appendix, Fig. S2A). Additionally, stratification by sex (female and male) showed no significant differences in hardness, elemental composition, or Raman spectra for incisors or molars (SI Appendix, Fig. S2B). Due to the distinct properties observed in enamel and dentin in PCA, separate multiple linear regression (MLR) analyses were performed to model predictors of hardness using five independent variables: EDS/Mg (at%), EDS/Fe (at%), EDS/P (at%), Ca/P, and C/P ratio (48). For enamel, the MLR model yielded an R-squared (R^2^) value of 0.8878, explaining approximately 88.78% of the variance in hardness and indicating a strong model fit (Fig. 5C). The ANOVA F-statistic was 38.00 (p < 0.0001), confirming the model’s statistical significance and indicating that the predictors explain a substantial portion of the variability in enamel hardness (SI Appendix, Table S2) (49). Specifically, atomic percentages of Mg, Fe, and the C/P ratio emerged as significant predictors. For each unit increase in Mg (at%) and Fe (at%), enamel hardness increased by 4.325 units (β_1_ = 4.325, p = 0.0282) and 5.721 units (β_2_ = 5.721, p = 0.0006), respectively. Conversely, the C/P ratio negatively predicted hardness (β_5_ = -1.358, p < 0.0001). It is worth noting that the C/P ratio reflects one peak for carbonate (1070 cm^−1^) and one for phosphate (960 cm^−1^). However, neither EDS/P alone nor the Ca/P ratio significantly influenced enamel hardness (β_3_ = -0.1153, p = 0.0698; β_4_ = 0.08087, p = 0.9325) (SI Appendix, Table S2).

The MLR model for dentin, though less robust, still reasonably explained hardness variance (R^2^ = 0.6683) (Fig. 5D), suggesting that additional factors may contribute to dentin hardness variability (50). Comparisons between dentin and enamel revealed that Mg (at%) was a significant predictor in both tissues, though its impact was stronger in dentin than enamel (SI Appendix, Table S3). Interestingly, Fe and the C/P ratio were highly significant predictors of enamel hardness, underscoring their greater influence on enamel hardness relative to dentin. This may reflect the role of iron in enamel mineralization (51, 52) and the substitution of carbonate for phosphate groups in hydroxyapatite crystals (53, 54).

## Discussion

Our results suggest that integrating mechanical and elemental analyses with imaging provides a deeper mechanistic understanding of enamel and dentin development. The differential roles of Fe and Mg, along with the non-linear relationship between Ca/P and hardness, suggest that mineralization involves complex crystal assembly processes beyond simple ion accumulation (55). Specifically, Fe’s contribution to enamel hardness and Mg’s role in dentin highlight tissue-specific mineralization strategies that may reflect distinct regulatory pathways, such as how collagen-ion (dentin) and amelogenin-ion (enamel) interactions may promote mineralization. Recent reports in Komodo dragon teeth have demonstrated the importance of Fe in the hardness of the hydroxyapatite enamel mineral (56), and our data supports this finding, indicating that Fe indeed contributes to hard tissue mechanical properties. Previous studies have reported that Mg content increases in the outer enamel layer due to the enamel-saliva ion exchange (57), however, because we used developing teeth, Mg access from the saliva seems improbable and may indeed suggest a biological role for Mg in hard tissue matrix development. Historically, compositional maturation has been associated with enhanced mechanical properties in mineralized tissues; our findings expand on this by quantifying the specific contributions of Mg and Fe to hardness (38). This suggests that increasing Fe content and reducing Mg could be effective strategies for improving hardness in synthetic enamel-like materials, potentially advancing biomaterial design and tissue engineering applications.

PCA analyses revealed distinct clustering patterns for enamel and dentin developmental stages. Enamel showed clear separation based on mineralization stages, particularly along PC1, while dentin clustered along PC2. This clustering pattern underscores the unique compositional changes in enamel during mineralization, contrasted with the more stable properties of dentin. Our findings showed that Mg content, in particular, distinguished enamel from dentin, supporting the hypothesis that Mg plays a key role in determining tissue-specific mineralization. This differentiation is consistent with our results showing stronger correlations between elemental composition and hardness in enamel than in dentin. In the context of the literature, previous studies have suggested that enamel and dentin develop through distinct pathways due to their roles in tooth function (13, 58). Our findings contribute to this understanding by identifying specific elemental markers that correlate with tissue type and mineralization stage. The significance of this clustering lies in advancing our ability to distinguish tissue types and developmental stages in complex mineralized tissues, which could aid in diagnostics and biomaterial design.

Through multiple linear regression (MLR), we identified Mg, Fe, and the C/P ratio as significant predictors of enamel hardness, with the model explaining 88.78% of the variance. In contrast, the MLR model for dentin was less predictive, indicating that additional factors likely contribute to its hardness. These results suggest that elemental composition is more directly tied to hardness in enamel than in dentin. Consistent with our PCA findings, enamel hardness varied significantly with changes in mineralization stages and elemental composition, while dentin remained relatively stable. These findings align with literature identifying Mg and Fe as influential in mineralization (59), where Fe is suggested to enhance enamel hardness by stabilizing crystal growth (52). This predictive model has notable implications for dental material development, offering a foundation for designing restorative materials that mimic the mechanical properties of natural enamel by targeting specific elemental compositions.

Future research should explore the mechanisms by which these ions are incorporated and contribute to tissue hardness, particularly through processes such as ion exchange and crystallization, specifically at the protein-ion level. Beyond extracted or exfoliated teeth, there is now an opportunity to investigate mineralization mechanisms dynamically in vivo or through developmental time points using various genetic mouse models. The role of Mg in both enamel and dentin raises questions about how ions are incorporated, reorganized, or replaced over time and how this might vary across different species or under pathological conditions (60, 61). Additionally, these findings could also provide valuable insights into other biological systems, such as vertebrate models with specialized dentition, archaeological samples to infer mineralization patterns in ancient populations, and genetic models where defects in mineralization pathways lead to dental and craniofacial mineralization defects. Expanding this research into other species and contexts would further elucidate the evolutionary and functional significance of tissue-specific mineralization strategies.

Overall, this study offers new insights into the mineralization dynamics and mechanical properties of developing craniofacial tissues, with particular emphasis on enamel, dentin, and bone. Our integrated approach highlights the value of combining multiple techniques to explore both composition and function. These findings open new avenues for future research, shedding light on mineralization mechanisms that could inform both biological and clinical applications. Together, our results suggest that mineral quality and composition drive tissue differentiation and could offer new biomarkers for studying hard tissue development across diverse biological systems.

## Materials and Methods

Animal experiments were approved and performed according to the IACUC panel of Children’s Hospital of Philadelphia. Mice were housed in a 12-hour light: dark cycle with food and water provided *ad libitum*. Mice were anesthetized with isoflurane and perfused with 1x PBS followed by 4% paraformaldehyde in PBS. Following careful degloving and decapitation, full skulls were further immersed and fixed in 4% paraformaldehyde for 24 hours on a shaker overnight at 4ºC. Samples were stored in 1x PBS at 4ºC.

### MicroCT

Acquisition details. A SCANCO MicroCT 45 machine (70kVP, 7.4um, 900ms) was used to scan full skulls in 70% ethanol. Following contouring of scans, DICOM files were exported. The incisors and first molar teeth were then segmented using BounTI (62)-with the following parameters: Initial Threshold – 10000, Target Threshold – 6000, Number of Iterations – 100 and Number of Segments – 9. The 9 segments were manually separated from the 10000 threshold segmentations to the 4 first molar teeth, the 4 incisors, and the rest of the bones and teeth as one segment. This was used as the seed to generate the segmentation. Lastly, the bones and other teeth were removed from the final segmentation, and a smoothing of 3 voxels was applied. To visualize the teeth of interest, volume renderings were produced in Avizo (Thermo Fisher Scientific, MA, USA) with greyscale (proxy for density) values coloured from blue to red, ranging from 5000 to 25000 in terms of the greyscale value. The segmentation regions were also overlaid on the 2-dimensional projections of the teeth with the same color scale as for the volume renderings. The volume and average grayscale value of each investigated tooth were recorded.

### Sample preparation

Following microCT, left-sided mandibles were collected and subsequently infiltrated with a graded series of ethanol for 15 minutes (50%, 70%, 90%, and 100%) followed by a 1:1 ethanol:resin mixture overnight. The samples were then embedded in 100% LR White resin (LR White Resin, Electron Microscopy Sciences, PA, USA) and polymerized at 65°C for 48 hours. After polymerization, the embedded specimens were sectioned using diamond blades (C.L. Sturkey, PA, USA) on a microtome (Leica, USA) to achieve a sagittal orientation that fully exposed both the incisor and the first molar. Then samples were polished using a surface grinder (PetroThin, Buehler, IL, USA). Prior to investigation, the sections were thoroughly rinsed with distilled water and dried.

### Scanning Electron microscopy (SEM) Imaging and Energy Dispersive X-ray Spectroscopy (EDS) Measurements

Electron microscope images were acquired using an FEI Quanta 600 FEG Mark II Environmental Scanning Electron Microscope (ESEM), which was operated in low vacuum mode at 0.38 Torr, an accelerating voltage of 15 kV, and a working distance of 10 mm. These settings were chosen to effectively identify and visualize regions for nanoindentation and subsequent compositional analysis. A total of six images were captured, with one image taken per sample. To determine the elemental composition of enamel and dentin, energy-dispersive X-ray spectroscopy (EDS) measurements were performed using an EDAX EDS detector integrated with the Quanta 600 ESEM without previous sputtering. The horizontal field width for all EDS measurements was set to 300 µm. Line scans of 100 µm in length (line width: 2 µm, resolution: 0.4 µm) were performed from enamel to dentin for each region on the incisors and molars, resulting in approximately 260 measuring points per line (Fig. 3A). For the bone regions, 75-µm line scans were used, with approximately 190 measuring points per scan.

### Nanoindentation

Mechanical characterization of rodent teeth and mandibles was conducted using an iMicro nanoindenter (Nanomechanics Inc., USA) housed at the University of Pennsylvania. The elastic modulus (E) and indentation hardness (H) were determined by indenting the specimens with a diamond Berkovich tip to a load of 50 mN at an indentation rate of

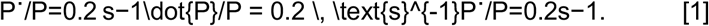

Continuous Stiffness Measurement (CSM) was employed to measure contact stiffness throughout the test (63). Specimens were mounted on aluminum pucks using Crystalbond 509, and the pucks were subsequently loaded into the nanoindenter’s sample tray for testing.

Nanoindentation tests were conducted on six distinct regions, including the secretory, transition, and maturation stages of the incisors, the cervical crown and cusp tip of the first molars, and the mandibular bone adjacent to the first molar. For each region on the incisors and molars, six measurement points were tested (Fig. 3A), while four points were tested on the mandibular bone. The distance between each measurement point was maintained at 30 μm.

### Raman Spectroscopy

Raman spectra were acquired using a Raman-Near Field Scanning Optical Microscopy (NSOM) system (NT-MDT Spectrum Instrument Ltd., Ireland) equipped with a 660-nm laser. The spectra were recorded over a spectral range of 295 to 1865 cm^−1^ with a spectral resolution of 1 cm^−1^, utilizing a diffraction grating of 600 lines/mm. Each Raman spectrum was corrected for fluorescence backgrounds by subtracting a seventh-order polynomial fitting curve using OMNIC 8 software (Thermo Fisher Scientific Inc., MA, USA). Following spectral acquisition, the mineral distribution was assessed based on the relative evaluation of specific band areas. The carbonate-to-phosphate ratio was determined by integrating the area under the bands associated with phosphate and carbonate components. Prior to the integration, all spectra underwent rubber band correction (RBC) with 15 iterations using OPUS 7.5 software (Bruker Optics, Ettlingen, Germany). After deconvolution, the integration regions were defined as ν_1_PO_4_^3−^ at 980–920 cm^−1^ and ν_3_CO_3_^2−^ at 1093–1057 cm^−1^. Peak parameters were obtained by fitting the 1030–1100 cm^−1^ region of the Raman spectrum using two Gaussian-Lorentzian curves, which were analyzed with OriginPro 2016b software (OriginLab Co., Northampton, MA, USA).

### Data Analysis

All results were expressed as the means ± standard deviation (SD). Statistical analysis was carried out using GraphPad Prism (Version 10.2.3, GraphPad Software, Boston, MA, USA) and Origin2021b (OriginLab, Northampton, MA, USA). To evaluate changes in enamel and dentin properties, either one-way ANOVA or mixed ANOVA with repeated measures was employed. One-way ANOVA was used to assess whether there were statistically significant differences in the means of measurements at specific developmental stages for each characterization method. Mixed ANOVA with repeated measures was applied when the position factor, as a within-subject variable, was included (e.g., for Raman microspectroscopy and EDS analyses). A significance level of p < 0.05 was used for all analyses.

### Principal Component Analysis and Multiple Linear Regression

Principal Component Analysis (PCA) was used to explore patterns and relationships among variables related to enamel and dentin properties. The analysis was conducted using GraphPad Prism on a dataset of six samples (three males, three females), including categorical variables (gender, tissue type, position) and continuous variables (NI-hardness, EDS/Mg (at%), EDS/Fe (at%), EDS/Ca (at%), EDS/P (at%), and C/P ratio). Continuous variables were standardized (mean = 0, SD = 1) to ensure comparability across scales. Principal components were selected based on parallel analysis using 1000 Monte Carlo simulations, retaining components with eigenvalues greater than those from the simulations at the 95% confidence level.

Multiple Linear Regression (MLR) was employed to evaluate how the selected variables (EDS/Mg (at%), EDS/Fe (at%), EDS/P (at%), Ca/P, and C/P ratio) influenced NI-hardness. The independent variables included the same continuous and categorical variables used in PCA. The analysis used the least squares method, assuming a Gaussian distribution of residuals. Multicollinearity was assessed using variance inflation factors (VIF), and model fit was evaluated through R-squared values. Diagnostic tests for normality (D’Agostino-Pearson, Anderson-Darling, Shapiro-Wilk, and Kolmogorov-Smirnov) were applied to the residuals.

## Acknowledgments

We would like to thank AZB & KDP for providing the inspiration for this story and for initiating the collaboration. We would also like to thank the lab members of the Goldsby Lab, Bhoj Lab, Ahrens-Nicklas Lab, and Vining Lab for their supportive and collegial environments where much of this work took place. Finally, we would like to thank our collaborators at the University of Pennsylvania Singh Center (Drs. Matthew Bruckman, Jaime Ford, Eric Stach), McKay Orthopaedic Research Laboratory PCMD MicroCT Core (Dr. Wen Sang), Dr. David L. Goldsby, and Dr. Ottman A. Tertuliano for their guidance and connections to key resources for this project. This work was partly carried out at the Singh Center for Nanotechnology, supported by the NSF National Nanotechnology Coordinated Infrastructure Program under grant NNCI-2025608.

Research reported in this publication was supported in part by a NIDCR Supplement to KK from the National Center for Advancing Translational Sciences of the National Institutes of Health under award number KL2TR001879.

## Notes

### Competing Interest Statement

The authors have declared no competing interest.

