## Supporting Information for "Multi-modal characterization of rodent tooth development"

### Supporting Information for
**Multi-modal characterization of rodent tooth development**

**Author List:**

Yuchen Jiang* [1], Kaitlin A. Katsura*[2,3,4], Nir Z. Badt[5], Marius Didziokas[6], Sonia Dougherty[2], David L. Goldsby[5], Elizabeth J. Bhoj[2,7 +], Kyle Vining[1,8 +]

*co-first

+co-senior/corresponding

Kyle Vining


**This PDF file includes:**

Supporting text
Figures S1 to S2
Tables S1 to S3

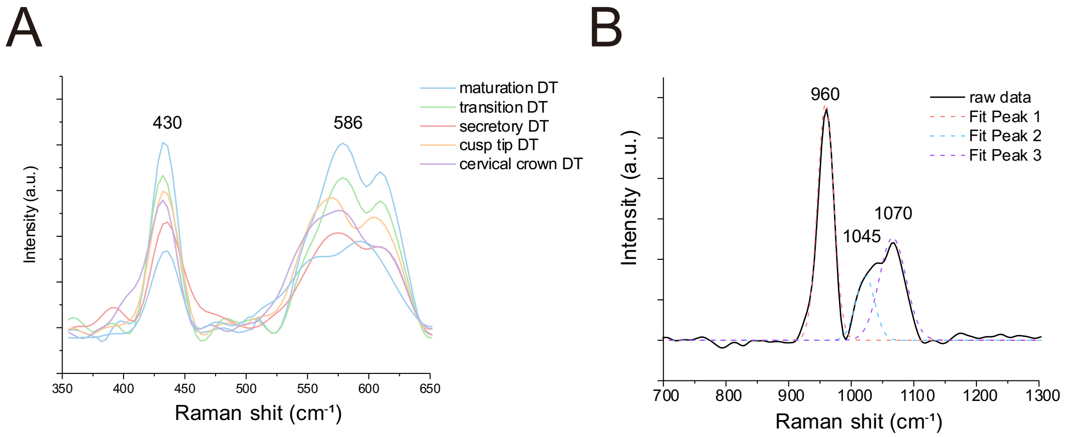


**Fig. S1.** (A) The spectral region of the $v_2$, $v_4$ phosphate group in dentin. (B) Raman sample fitting for peak area integration. Three Gaussian-Lorentzian curves were fitted to obtain the peak area for C/P ratio calculation.

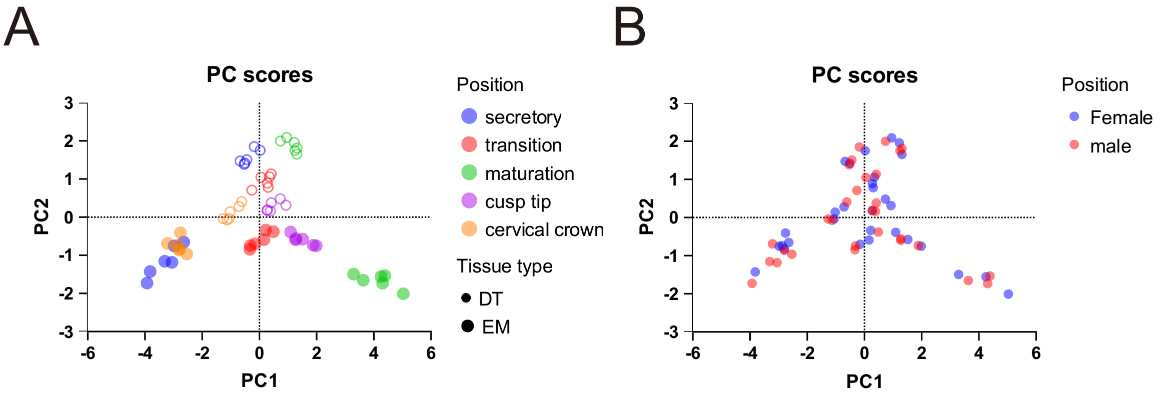


**Fig. S2.** Biplots for PCA analysis. (A) PCA biplot showing the clustering of data points specified with position and tissue type in incisor and molar based on PC1 and PC2. (B) Another biplot demonstrates that there are no differences in incisor or molars corresponding to hardness, elemental and chemical compositions.

**Table S1.** Tabular results for PC selection, where the first two principal components (PC1 and PC2) capture the largest variance (76.63%) in the dataset. The table also shows the eigenvalues for each (PC), along with the explained variance percentage and the cumulative variance explained by these components.

| Table Analyzed | PCA | | | | | | |
|---|---|---|---|---|---|---|---|
| PC summary | PC1 | PC2 | PC3 | PC4 | PC5 | PC6 | PC7 |
| Eigenvalue | 4.269 | 1.290 | 0.7860 | 0.4257 | 0.1397 | 0.05861 | 0.03130 |
| Proportion of variance | 60.99% | 18.42% | 11.23% | 6.08% | 2.00% | 0.84% | 0.45% |
| Cumulative proportion of variance | 60.99% | 79.41% | 90.64% | 96.72% | 98.72% | 99.55% | 100.00% |
| Component selection | Selected | Selected | | | | | |

**Table S2.** Tabular results of multiple linear regression for enamel hardness. Multiple Linear Regression (MLR) results summarizing the sum of squares (SS), degree of freedoms (DF), mean squares (MS), F-statistics (F), standard errors, and p-values for the predictor variables across input variables. The rows correspond to the independent variables, while the columns display the statistical outputs from the regression analysis. Significant predictors, including Fe (at%), C/P and Mg (at%), are identified by their low p-values, indicating their contribution to the model.

| **Model Analysis of Variance** | **SS** | **DF** | **MS** | **F (DFn, DFd)** | **P value** |
|---|---|---|---|---|---|
| Regression | 11.35 | 5 | 2.269 | F (5, 24) = 38.00 | P<0.0001 |
| EDS/Mg (at%) | 0.3258 | 1 | 0.3258 | F (1, 24) = 5.456 | P=0.0282 |
| EDS/Fe (at%) | 0.9433 | 1 | 0.9433 | F (1, 24) = 15.79 | P=0.0006 |
| EDS/P (at%) | 0.2151 | 1 | 0.2151 | F (1, 24) = 3.602 | P=0.0698 |
| Ca/P | 0.0004378 | 1 | 0.0004378 | F (1, 24) = 0.007330 | P=0.9325 |
| C/P | 1.451 | 1 | 1.451 | F (1, 24) = 24.30 | P<0.0001 |
| Residual | 1.433 | 24 | 0.05973 | | |
| Total | 12.78 | 29 | | | |
| **Parameter estimates** | **Variable** | **Estimate** | **Standard error** | **P value** | **P value summary** |
| β0 | Intercept | 1.004 | 1.977 | 0.6163 | ns |
| β1 | EDS/Mg (at%) | 4.325 | 1.852 | 0.0282 | * |
| β2 | EDS/Fe (at%) | 5.721 | 1.440 | 0.0006 | *** |
| β3 | EDS/P (at%) | -0.1153 | 0.06072 | 0.0698 | ns |
| β4 | Ca/P | 0.08087 | 0.9446 | 0.9325 | ns |
| β5 | C/P | -1.358 | 0.2756 | <0.0001 | **** |

**Table S3.** Tabular results of multiple linear regression for dentin hardness. Mg (at%) is the significant predictor identified by their low p-values, indicating their contribution to the model.

| **Analysis of Variance** | **SS** | **DF** | **MS** | **F (DFn, DFd)** | **P value** |
|---|---|---|---|---|---|
| Regression | 0.2909 | 5 | 0.05818 | F (5, 24) = 9.673 | P<0.0001 |
| EDS/Mg (at%) | 0.2085 | 1 | 0.2085 | F (1, 24) = 34.67 | P<0.0001 |
| EDS/Fe (at%) | 0.003770 | 1 | 0.003770 | F (1, 24) = 0.6268 | P=0.4363 |
| EDS/P (at%) | 0.003989 | 1 | 0.003989 | F (1, 24) = 0.6632 | P=0.4234 |
| Ca/P | 0.01365 | 1 | 0.01365 | F (1, 24) = 2.269 | P=0.1451 |
| C/P | 0.0008641 | 1 | 0.0008641 | F (1, 24) = 0.1437 | P=0.7080 |
| Residual | 0.1444 | 24 | 0.006015 | | |
| Total | 0.4353 | 29 | | | |
| **Parameter estimates** | **Variable** | **Estimate** | **Standard error** | **P value** | **P value summary** |
| β0 | Intercept | -0.6093 | 0.7692 | 0.4361 | ns |
| β1 | EDS/Mg (at%) | 1.164 | 0.1978 | <0.0001 | **** |
| β2 | EDS/Fe (at%) | -0.4389 | 0.5544 | 0.4363 | ns |
| β3 | EDS/P (at%) | 0.03362 | 0.04128 | 0.4234 | ns |
| β4 | Ca/P | 0.3374 | 0.2240 | 0.1451 | ns |
| β5 | C/P | 0.01299 | 0.03428 | 0.7080 | ns |
